# A spatial fingerprint of land-water linkage of biodiversity uncovered by remote sensing and environmental DNA

**DOI:** 10.1101/2021.10.27.466050

**Authors:** Heng Zhang, Elvira Mächler, Felix Morsdorf, Pascal A. Niklaus, Michael E. Schaepman, Florian Altermatt

## Abstract

Aquatic and terrestrial ecosystems are tightly connected via spatial flows of organisms and resources. Such land-water linkages integrate biodiversity across ecosystems and suggest a spatial association of aquatic and terrestrial biodiversity. However, knowledge about this spatial extent is limited. By combining satellite remote sensing (RS) and environmental DNA (eDNA) extraction from river water across a 740-km^2^ mountainous catchment, we identify a characteristic spatial land-water fingerprint. Specifically, we find a spatial association of riverine eDNA diversity with RS spectral diversity of terrestrial ecosystems upstream, peaking at a 400 m distance yet still detectable up to a 3.3 km radius. Our findings testify that biodiversity patterns in rivers can be linked to the functional diversity of surrounding terrestrial ecosystems and provide a dominant scale at which these linkages are strongest. Such spatially explicit information is necessary for a functional understanding of land-water linkages and provides a reference scale for adequate conservation and landscape management decisions.

## 1. Introduction

Understanding the spatial distribution of biodiversity and its linkage across ecosystem types is essential, especially in an era of increasing human modifications of natural landscapes ^1,2^. It is well-established that species and ecosystem functional diversity are unevenly distributed across landscapes, with pronounced diversity hot and cold spots ^3,4^. Extensive work has also demonstrated how ecosystems more diverse in species are more productive and stable ^5–7^. Intriguingly, however, most past work has focused on individual ecosystem types, such as forests, grasslands, or aquatic ecosystems, thereby neglecting a possible co-variation of biodiversity across different ecosystems ^8^. Indeed, only very recently the relevance of spatial scaling of biodiversity and ecosystem functioning research and the dependence on the spatial extent has been postulated ^9,10^

Natural ecosystems, and the biodiversity therein, are often linked to each other through flows of organisms and resources ^11,12^. One of the most prominent examples is the coupling of aquatic to terrestrial ecosystems ^13,14^. Aquatic ecosystems are not only highly biodiverse yet threatened by anthropogenic activities ^15,16^, but also strongly interlinked with surrounding terrestrial ecosystems through the characteristic fractal structure of riverine networks across most landscapes worldwide ^17,18^. Consequently, in these systems, the interaction of one ecosystem resulting in an imprint on the diversity of the other ecosystem is expected, with implications for land management and conservation. Nevertheless, little is known about the occurrence and extent of such imprints, particularly regarding the spatial range at which such an interaction modulates local biodiversity.

To assess possible spatial linkages of diversity across ecosystem types, biodiversity must be quantified in scalable manners. Classically, biodiversity is directly quantified by counting individual species, for example, through inventories conducted along transects or in plots of defined size. This approach, however, is inherently limited for spatial upscaling and cross-ecosystem comparisons ^9^. Currently, two recent technological advances are revolutionizing biodiversity sciences, overcoming limitations with taxonomic and functional coverage, and the possibility to be spatially scaled. The first advancement is through remote sensing (RS) methods, which use portable, airborne, or satellite devices to characterize the ecosystem structurally, taxonomically, or physiologically by measuring reflected or emitted radiation at a distance ^19–21^. RS is particularly capable of characterizing terrestrial plant communities and a prime method for measuring essential biodiversity variables (EBVs) ^19–21^. Particularly, RS can map terrestrial ecosystem functional traits and diversity at regional to global scales with resolutions down to a meter, enabling the upscaling of biodiversity from local composition to ecosystem levels ^22–25^. The second advancement is through environmental DNA (eDNA) metabarcoding, which uses DNA extracted from environmental samples to quantify biodiversity across the tree of life ^26–30^. eDNA metabarcoding is widely used in aquatic ecosystem studies, where it is becoming a standard for biodiversity assessments ^31–36^. The passive transport of DNA in water makes it a particularly efficient method in riverine systems, as the flow along the riverine network carries and integrates biodiversity information over the catchment ^37–40^, and can be used for estimating spatial patterns of biodiversity at the landscape level ^41,42^.

Essentially, RS and eDNA metabarcoding complement each other in biodiversity detection. eDNA can detect bacteria, invertebrates, and vertebrates that are largely inaccessible for RS, while RS can monitor ecosystem physiological and structural diversity impossible to draw from eDNA data. Therefore, a combination of RS and eDNA can provide a holistic view of biodiversity for isolated and mosaicked ecosystems ^43,44^ and allows to uncover land-water linkages of biodiversity at the landscape level ^45,46^.

Here, we quantified the spatial extent of a linkage of biodiversity between aquatic and terrestrial ecosystems by combining eDNA sampling and RS in a 740-km^2^ river drainage basin. We assessed aquatic biodiversity along the river network using eDNA and matched it to terrestrial ecosystem functional diversity in the catchment based on Sentinel-2 Multi-Spectral Instrument (MSI) satellite data. Specifically, we identified the spatial range within which the functional diversity of the terrestrial vegetation was associated with the taxonomic diversity in the riverine ecosystems and determined at what spatial scale this linkage was the highest. Thereby, combining eDNA and RS, we provide a first spatially explicit integration of land-water linkage of biodiversity, and identify a characteristic spatial fingerprint across aquatic-terrestrial ecosystem boundaries at the landscape level.

## 2. Results

We combined assessments of aquatic biodiversity using eDNA and terrestrial diversity based on Sentinel-2 Multi-Spectral Instrument (MSI) satellite data in the 740 km^2^ river Thur catchment (Fig. 1). The river Thur catchment is located in the northeastern part of Switzerland. It covers a mountainous landscape with an elevation gradient ranging from 460 m to 2423 m a.s.l. and contains a mosaicked landscape of urban, agricultural and forested terrestrial ecosystem types.

**Fig. 1.**
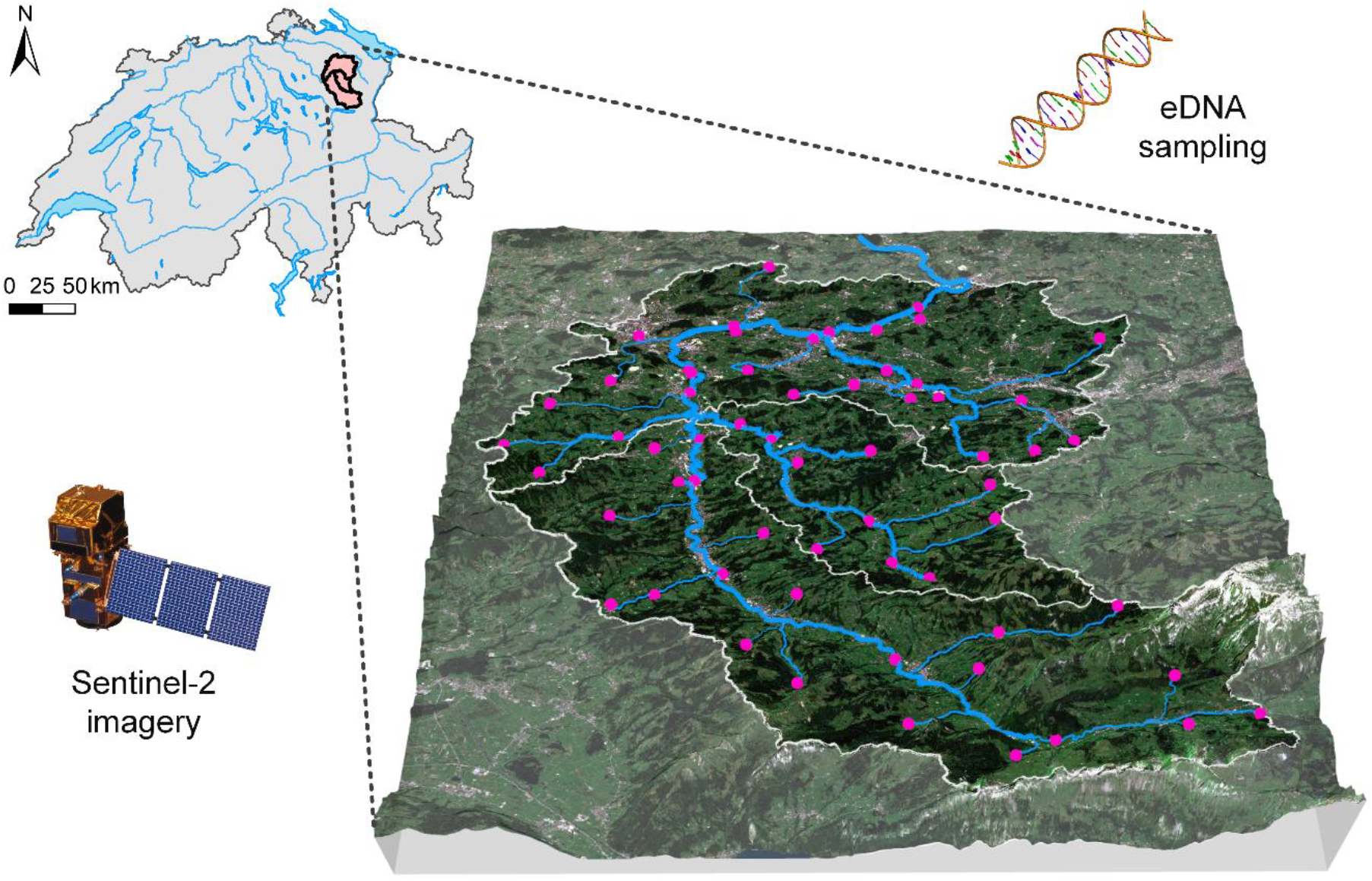
Location of the Thur river catchment in Switzerland and eDNA sampling sites. Pink dots are 61 eDNA sampling sites. The blue lines represent river channels draining in a Northward direction. White lines indicate the boundaries of the main catchment and its three subcatchments (Thur, Glatt, and Necker).

### eDNA-derived biodiversity in aquatic ecosystem

We conducted eDNA water sampling at 61 sites along the river network, representatively covering the whole 740 km^2^ catchment (Fig. 1). All water samples were filtered, the DNA extracted, and sequenced using generic COI-specific primers targeting a broad range of pro- and eukaryotic organisms. Detailed procedures are described in the Methods section and in Mächler *et al.*, 2019 and 2021 ^47,48^. We received a total of 26,519,031 reads that were clustered into 10,962 zero-radius operational taxonomic units (ZOTUs) with 2404 ± 216 (mean ± standard error) number of reads per ZOTU as a proxy of taxonomic diversity.

To describe different aspects of biodiversity across all eDNA samples, we used Hill numbers, which are a compatible statistical framework considering both occurrence and abundance information ^48–51^. In this framework, the evenness of biodiversity patterns gets more weight with increasing Hill number q orders. Here, we calculated Hill numbers with order q = 0, 1, and 2, which correspond to species richness, the exponential of Shannon diversity, and the inverse of the Simpson index, respectively (see Methods, Fig. 2), after removing very rare ZOTUs (occurrence < 0.005% in total, see details in Methods section). We observed strong and highly uneven biodiversity patterns across the catchment, with a strong and significant positive correlation between biodiversity and Strahler order (Fig. S1 a-c; p-value < 0.05), and a decreasing trend of biodiversity at increasing elevation (Fig. S1 d-f).

**Fig. 2.**
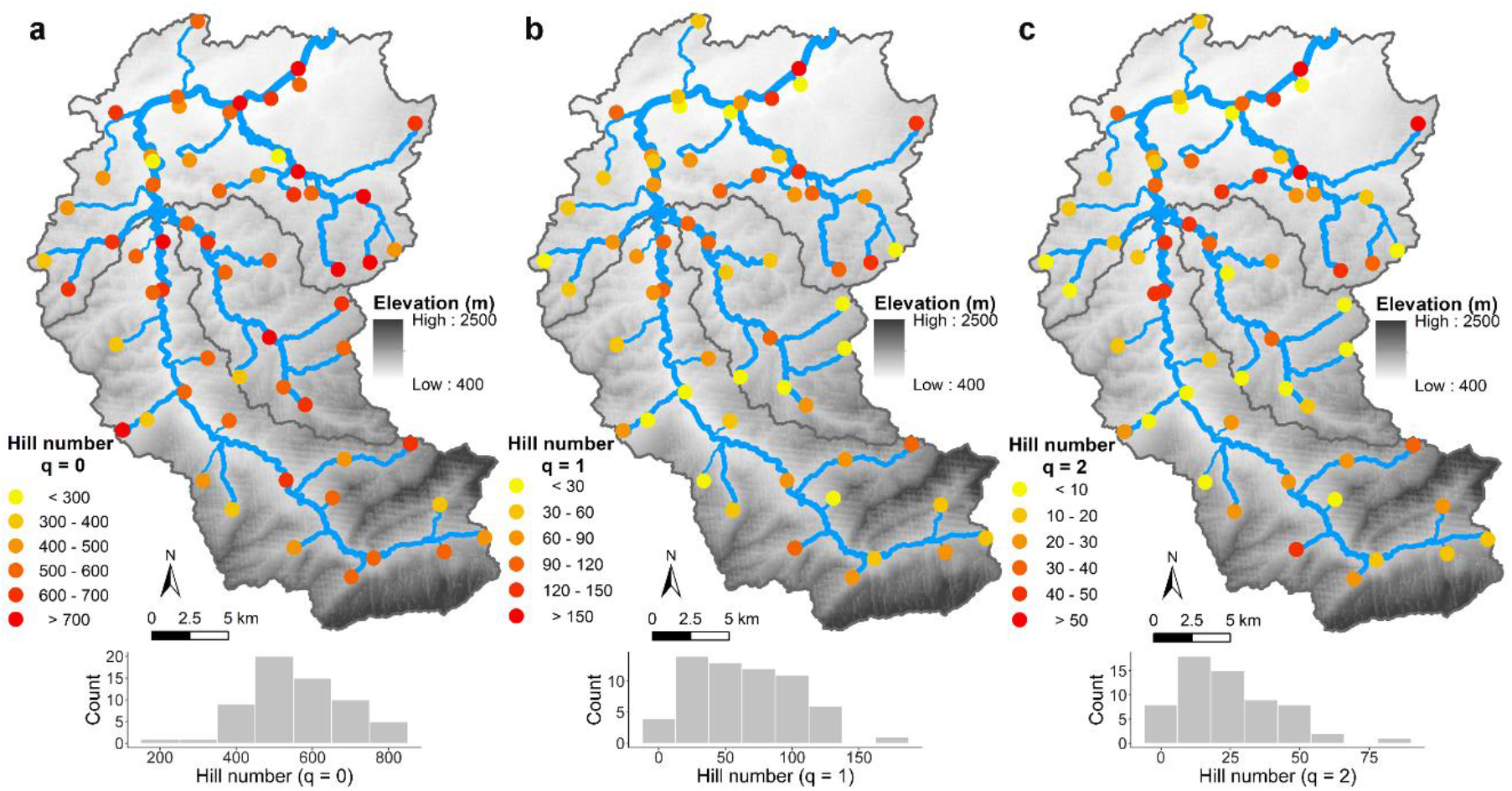
Distribution of biodiversity in Thur river catchment. Hill numbers were used to describe biodiversity of eDNA samples in the river network. Spatial patterns and histograms on distribution of diversity using Hill numbers with order **a** q = 0, **b** q = 1, and **c** q = 2 are given. They correspond to species richness (order q = 0), the exponential of Shannon diversity (order q = 1), and the Simpson index (order q = 2), respectively.

### RS-derived physiological traits and functional diversity in terrestrial ecosystems

We adapted a spatially continuous method, which was generalized to Sentinel-2 MSI satellite data, to map the terrestrial ecosystem functional diversity (a metric in EBVs) at a 20 × 20 m resolution ^22,52,53^. We used chlorophyll content (CHL), anthocyanin content (ANT), carotenoid content (CAR), and water content (WAT) to represent four physiological trait dimensions (see Methods) as direct proxies of functional diversity (Fig. 3). These spectral components capture plant physiological traits that integrate different components of terrestrial ecosystem functions, and thus functional diversity, related to the presence and conditions of vegetation ^52,54^.

**Fig. 3.**
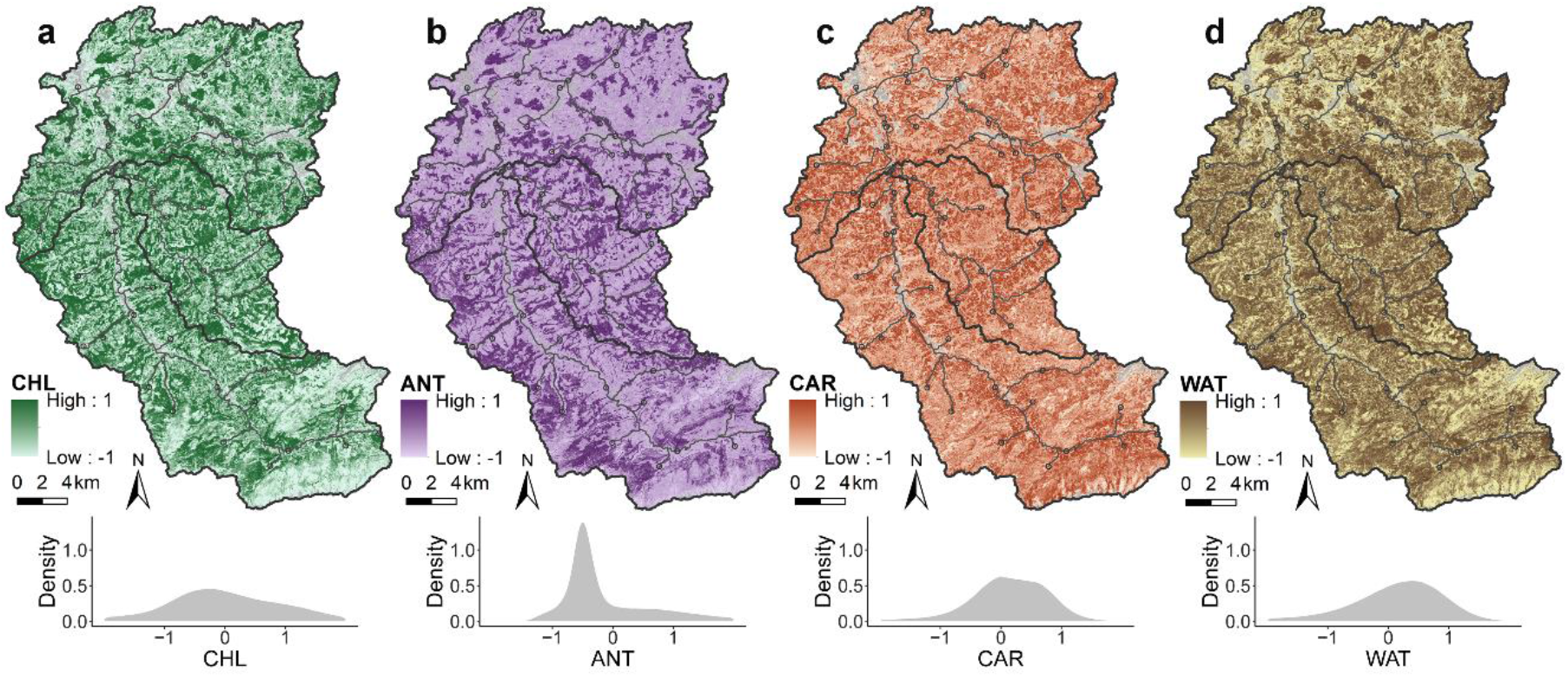
Functional trait diversity assessed through physiological trait characteristics of the terrestrial landscape in the Thur river catchment. **a** Map and density plot of chlorophyll content (CHL), represented by the red-edge chlorophyll index (*CI*_re_). **b** Map and density plot of anthocyanin content (ANT), represented by the anthocyanin reflectance index 1 (*ARI1*). **c** Map and density plot of carotenoid content (CAR), represented by the plant senescence reflectance index (*PSRI*). **d** Map and density plot of water content (WAT), represented by the normalized difference infrared index (*NDII*). Non-vegetated pixels were masked out (grey area), and all traits were normalized.

For each sampling site, we produced a catchment map based on the digital elevation model (DEM) and created distance buffers with spatial intervals, within which functional divergence (FDiv) was calculated (Fig. 4, see Methods section). The mean value of FDiv (± standard deviation (SD)) across distance is 0.665 (± 0.025). As the distance increased, the range of FDiv dropped from 0.216 (distance = 50 m) to 0.049 (distance = 20 km).

**Fig. 4.**
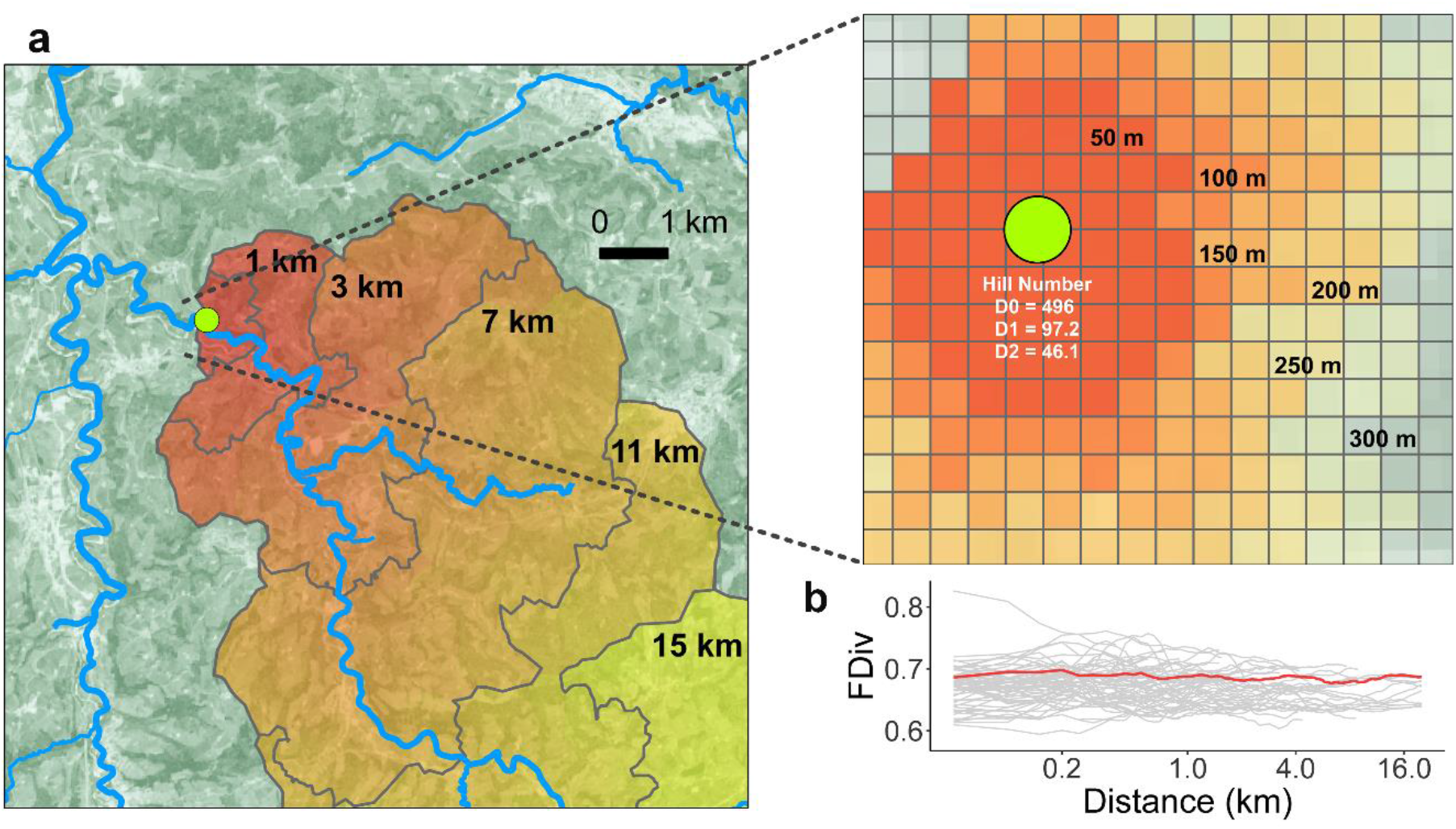
Spatial distribution of terrestrial ecosystem functional diversity based on catchment and distance buffers of the eDNA sampling site. **a** Catchment map with distance buffers of site No. 28 as an example. The spatial interval is 0.05 km for 0–10 km and 0.1 km for 10–20 km. **b** Functional divergence (FDiv) with upstream distance given for 61 eDNA sampling sites (grey lines; the example site No. 28 is highlighted as red line). We calculated FDiv by collecting four-dimensional trait value vectors from pixels covered by the distance buffer (for details and equations, see Methods section). Non-vegetated pixels were masked out before computation.

### The land-water linkage of biodiversity

We employed a model II simple linear regression to assess the association between eDNA-derived biodiversity (Hill numbers) and the RS-derived terrestrial ecosystem functional diversity (FDiv) across distance, using R^2^ as the goodness of fit. Uncertainties were estimated by a bootstrap framework (see Methods for details).

The linear regression analysis reveals a unimodal association between the eDNA-based (aquatic) Hill numbers and the RS-based (terrestrial) FDiv as the upstream distance to sampling sites increases, with a linkage signal of up to 3.3 km radius upstream (Fig. 5). The distances with the highest R^2^ (distance with maximal land-water imprint) vary across orders of q. For q = 0, this distance with the strongest imprint is 400 m; for q = 1, it is 350 and 800 m, respectively; for q = 2, it is 350 and 850 m, respectively. The strong effect of ZOTU-level richness decreases with increasing Hill number (Fig. 5), suggesting that the rare taxa contribute most to the observed land-water linkage. Possibly, this could be ascribed to the decreasing contributions from the less abundant taxonomic groups after increasing the weight of abundance (increasing Hill number order q), as an abundant taxonomic group may swamp the effect of the less abundant ones. In addition, it highlights the importance of rare taxa contributing to overall beta-diversity ^55^ and the negative effect of large-scale homogenization of biodiversity ^56^, which results not only in an erosion of beta-diversity within one ecosystem but has also a cascading negative effect on other ecosystems.

**Fig. 5.**
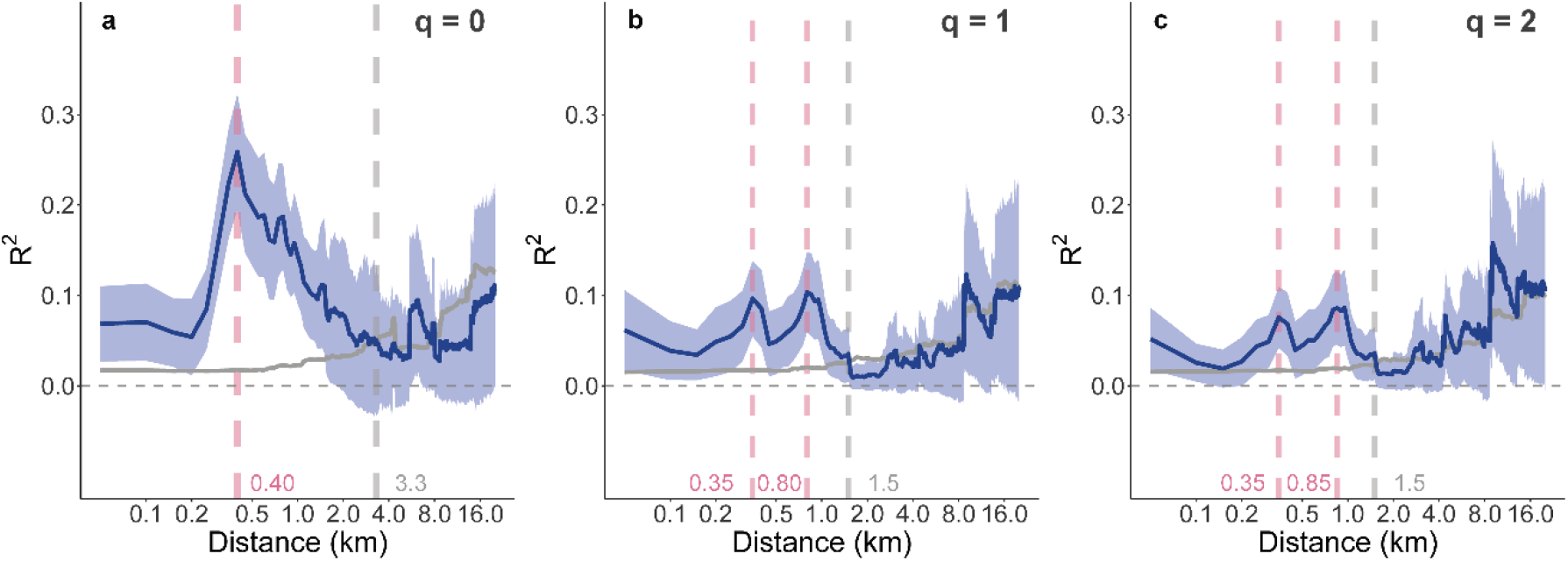
Association between eDNA-derived biodiversity assessed in the river water and RS-derived terrestrial ecosystem functional diversity across increasing upstream distance in the Thur river catchment. The R^2^ of the linear regression (± standard deviation, blue lines) between eDNA-based Hill numbers with order **a** q = 0, **b** q = 1, and **c** q = 2, and RS-based functional divergence (FDiv) across distance are given. The R^2^ of the null models is shown in grey lines.

We developed null models to corroborate the robustness of the observed spatial extent of the land-water linkage, by randomly shuffling the locations of all pixels within the river catchment. Then, we assessed whether and at what spatial extent such a land-water linkage of biodiversity exists in a null-model scenario (see Methods). We found that the R^2^ of our sampling was always greater than the null model for distances < 3.3 km for q = 0, < 1.5 km for both q = 1 and 2, respectively (Fig. 5). These results testify that biodiversity in riverine ecosystems can be linked to the functional diversity of surrounding terrestrial ecosystems, with the strongest association occurring at a spatial extent of several hundred meters.

To disentangle the observed land-water linkage of biodiversity, we mapped the ZOTUs against a customized MIDORI Reference 2 database for taxonomic information, which allowed us to identify the taxonomic affiliation of the most prominent ZOTUs and read numbers at phylum and class level, respectively (Fig. 6). Abundant affiliations both with respect to ZOTU richness and read numbers were found for Arthropods (especially Insecta), Ascomycota (a fungi phylum), and Bacillariophyta (i.e., diatoms), and ZOTUs across all groups originated from organisms inhabiting both aquatic and terrestrial environments (Fig. 6). We subsampled the eDNA data based on the taxonomic information to evaluate individual contributions across major taxonomic groups. Specifically, we calculated the relative abundances at the phylum level and assessed their associations with FDiv across distance. Among all the major taxonomic groups, we detected strong associations in Bacillariophyta, Chordata, Ascomycota, Cnidaria, Rotifera, Amoebozoa, Chlorophyta, Cryptophyta, and Porifera, although the spatial extents were varying (Fig. S2). Importantly, these results show that the land-water linkage of biodiversity included contributions of aquatic and terrestrial origins, thus reflecting both an integrated signal of biodiversity across ecosystems and a signal of local ecosystem biodiversity.

**Fig. 6.**
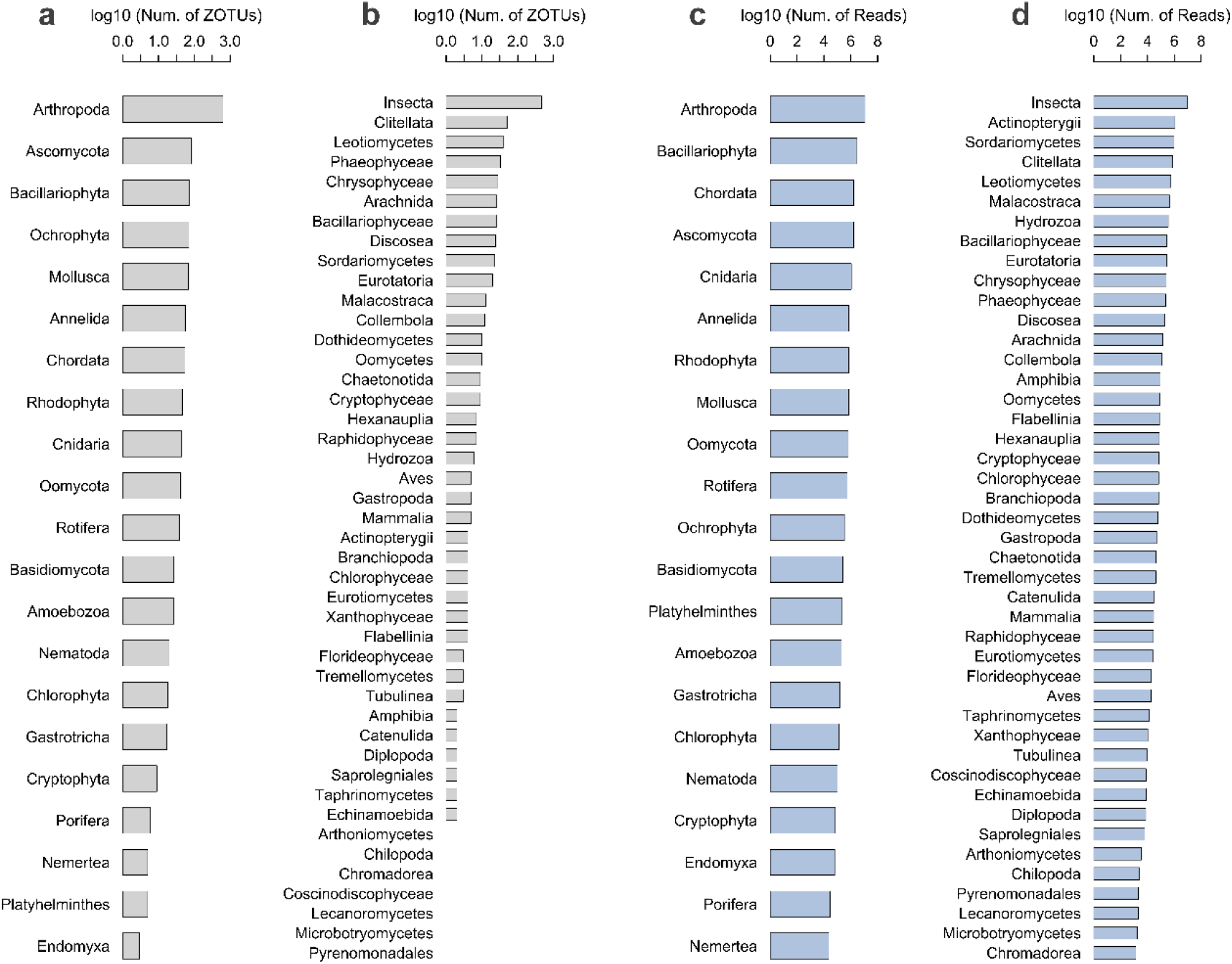
Number of ZOTUs and reads in the eDNA data. **a** Number of ZOTUs at the phylum level. **b** Number of ZOTUs at the class level. **c** Number of reads at the phylum level. **d** Number of reads at the class level. ZOTUs with occurrences less than three at the phylum level were removed to avoid spurious effects. All numbers were log_10_-transformed before plotting. The taxonomic information of eDNA data indicates a combination of aquatic and terrestrial origins.

## 3. Discussion

Combining eDNA sampling and multispectral remote sensing imagery (Fig. 1), we demonstrated a spatial association of biodiversity between aquatic and terrestrial ecosystems and gave a spatially explicit quantification of its peak strength, peaking across a catchment section at a 400 m radius upstream around the aquatic sampling site (Fig. 5). Overall, the unimodal signal of the land-water linkage of biodiversity covers a range of up to 3.3 km upstream, indicating that a place in a river and surrounding terrestrial ecosystems are closely interlinked, with a tight connection in terms of biodiversity. Furthermore, for the first time, we provide a specific and scalable approach to quantify the spatial extent of such linkages across ecosystems types and identify a characteristic spatial land-water fingerprint.

The characterization of the terrestrial ecosystems from a biodiversity perspective was based on multiple physiological trait dimensions (Fig. 3), capturing major components of the dominant vegetation cover. Contrary to traditional biodiversity surveys and estimates, which are often limited to small scales and numbers of sites and depend on specific taxonomic knowledge, our approach using high-resolution satellite RS data is not only capable of depicting regional and spatially continuous characteristics of biodiversity, but can be directly applied and scaled to map terrestrial biodiversity across all river catchments worldwide. Additionally, the characterization of aquatic biodiversity using eDNA allows a scaling across space and time, and most importantly, does not depend on prior knowledge on the occurrence of specific taxa. Thereby, this eDNA and RS combination approach could contribute to a global understanding of biodiversity patterns. Our method can, in principle, be applied and transferred to all land-water ecosystems worldwide, and may be especially useful to uncover biodiversity patterns in understudied regions, such as regions beyond Europe and North America.

In this study, we identified a strong fingerprint of land-water linkage of biodiversity, with a metric of terrestrial ecosystem functional diversity developed on a combination of four physiological trait components of vegetation. These four physiological traits are proved to be able to capture major ecosystem functions of vegetation ^52^. To evaluate the relative individual importance of these components, namely CHL, CAR, ANT, and WAT, we removed one dimension at each time and repeated the calculation process. We found that the maximum values of R^2^ dropped remarkably when CHL or WAT was removed (Fig. S3 & Tab. S1). Moreover, the unimodal shape was flatter after both CHL and WAT were removed (Fig. S4 & Tab. S1). These indicate that CHL and WAT, inherently representing the photosynthesis activity of vegetation and thus a proxy of productivity, mainly characterize the spatial fingerprint of land-water linkage of biodiversity.

For the characterization of the aquatic biodiversity (Fig. 2), we used a generic COI marker amplifying eDNA signals across a wide range of taxa, yet predominantly used to target invertebrates. Although a large proportion of retrieved sequences aligned with macro-and micro-invertebrates, we covered a wider breadth of taxa regarding ZOTUs, including microbes and vertebrates. Because the coverage of these organisms is highly variable in the respective reference databases ^57^, we applied a taxonomy-free approach using ZOTUs only to not depend on such databases. This approach covers a broader taxonomic breadth yet does not address the contribution of individual taxonomic groups. Still, according to the taxonomic information of our eDNA data (Fig. 6), we observed that ZOTUs originated from aquatic and terrestrial environments both contributed to the land-water linkage of biodiversity. Then, we also evaluated the relative contribution of each of the major taxonomic groups at the phylum level to the spatial land-water fingerprint by omitting one of these major taxonomic groups at a time and repeating the calculations. Intriguingly, the association pattern was almost the same regardless of which taxonomic group was omitted (Fig. S5), suggesting that the land-water fingerprint of biodiversity is highly robust and thus does not depend on a single major organismal group.

Importantly, the unimodal shape of the linkage of biodiversity was not caused by variations of vegetation productivity, suggesting that the heterogeneity and not the productivity of terrestrial ecosystems contributes to local aquatic biodiversity. We tested this by firstly calculating the enhanced vegetation index (EVI) to represent vegetation productivity ^58^. Then, we adopted type I ANOVA tests to evaluate the relative contributions of EVI and FDiv to the Hill numbers across distance (see Methods; Fig. S6). In addition, we also found that EVI and FDiv were not correlated at distances < 8.0 km (Fig. S7). Together, this evidences that the unimodal signal of land-water linkage of biodiversity cannot be ascribed to variations of vegetation productivity.

The methodology to assess the spatial fingerprint of land-water linkage of biodiversity proved to be an efficient way to uncover an underlying picture of biodiversity in spatially coupled ecosystems, by combining *in situ* measures of eDNA and regional data of RS. Both eDNA metabarcoding and RS are capable of assessing biodiversity across scales because of easy access to vast quantities of information with high robustness and accuracy, non-invasive and standardized procedures, and relatively low costs ^59–63^. Therefore, the methods applied here can contribute to next-generation biodiversity monitoring at regional to global scales ^64^.

The spatial fingerprint of land-water linkage of biodiversity detected is robust and may be even more resolved when the spatio-temporal matching of the two approaches is increased. Our study adopted Sentinel-2 MSI Level-2A bottom of atmosphere reflectance for RS measurements. It was generated on Level-1C top of atmosphere reflectance and is less affected by clouds or aerosols. Therefore, it is more accurate in mapping the physiological traits of vegetation. Due to the lack of Level-2A reflectance in 2016, we used Level-2A reflectance in 2017 for calculation in order to match the eDNA sampling at the respective seasonal time point. While there is likely seasonality in both RS and eDNA data ^65,66^, the inter-annual variation in RS between 2016 and 2017 is relatively minor, being testified by a very high correlation of corresponding bands and physiological trait indices on Level-1C data between 2016 and 2017 (Tab. S2 & S3). Additionally, the meteorological conditions were very similar between 2016 and 2017, and both years were close to the normal condition in terms of temperature and precipitation (Tab. S4). Thus, the spatial fingerprint is robust across years, at least when the land cover and meteorological conditions are not changing. In reverse, the method may be directly applicable to detecting land-use changes, as a change in the magnitude and extent of the spatial fingerprint may be expected.

In conclusion, we uncovered a spatially explicit land-water linkage of biodiversity in a large mountainous catchment by using eDNA sampling and satellite remote sensing imagery. The linkage of biodiversity between rivers and surrounding terrestrial landscapes covers a section in the catchment with a radius of around 3 km, with a maximum at 400 m, identifying a characteristic fingerprint of land-water linkage of biodiversity in spatially coupled ecosystems. While developed in a mountainous region with different major land cover types, including forest, grassland, agriculture, and urban areas, our method does not depend on specific organismal groups, thus can be used for all regions with mosaicked land cover types, providing a globally applicable basis for biodiversity conservation and land management.

## 4. Methods

### eDNA sampling in the Thur river network

The Thur catchment covers an area of 740 km^2^ with three main river branches (Thur, Glatt, and Necker) and the main land covers including forest (29.0%), arable and grassland (56.0%), urban area (10.2%), unproductive land (3.6%), and water (1.2%) land types (data from Swiss Federal Statistical Office, 2015. website: https://www.bfs.admin.ch/bfs/en/home/services/geostat/swiss-federal-statistics-geodata/land-use-cover-suitability/swiss-land-use-statistics/land-use.html). A systematic eDNA sampling was conducted in June 2016 under base-flow conditions. The detailed sampling, laboratory work, and subsequent bioinformatic analyses are described in Mächler *et al.*, 2019 and 2021 ^47,48^, who mostly analyzed the dataset with respect to the diversity of a small subset of all organisms and methodological details of the eDNA sampling, respectively. In total, we collected 183 water samples at 61 sites (three individual replicates per site) in the dendritic river network. For each replicate, 250 ml of river water was filtered on site using GF/F filters (pore size 0.7 um Whatman International Ltd.), and the filters were then immediately stored at −20 °C. Subsequently, DNA was extracted in a specifically dedicated clean lab, using the DNeasy Blood and Tissue Kit (Qiagen GmbH). Handling and extraction of all replicates were done in a randomized order. We performed two PCR runs with the Illumina MiSeq dual-barcoded two-step PCR amplicon sequencing protocol by targeting a short barcode region of the cytochrome c oxidase I (COI) ^67^. We used primers containing an Illumina adaptor-specific tail, a heterogeneity spacer, and the amplicon target site in the first run, and the Nextera XT Index Kit v2 for indexing in the second run. Filter controls (FC), extraction controls (EC), positive and negative PCR controls (PC, NC) were run alongside. The sequence data were subsequently demultiplexed, and the quality of the reads was checked with FastQC ^68^. Then, we end-trimmed (usearch, version 10.0.240), merged the raw reads (Flash, version 1.2.11), removed primer sites (cutadapt, version 1.12), and quality-filtered the data (prinseq-lite, version 0.20.4). Next, we used UNOISE3 (usearch, version 10.0.240) to determine ZOTUs, and performed an additional clustering at 99% sequence identity to reduce sequence diversity. Before final use, the resulting ZOTUs were checked for stop codons with invertebrate mitochondrial code, and to only contain an intact open reading frame.

We merged the ZOTU abundances of the three replicates at each site and got 26,519,031 reads clustered into 10,962 ZOTUs. Then, we calculated the relative abundance for each ZOTU at all sampling sites. To alleviate uncertainties, we filtered out the ZOTUs with less than 0.005% occurrence in total (i.e., <1326 total reads) and finally used 24,471,930 reads clustered into 1,394 ZOTUs for all analyses. Taxonomic information at the phylum and the class level for all ZOTUs was acquired by mapping against a customized MIDORI Reference 2 (UNIQ/GB242) database. After that, we computed relative abundance for each ZOTU at each site, subsequently referred to as our eDNA data.

### Hill numbers as metrics of eDNA-derived biodiversity

We used Hill numbers as a scalable metric to describe eDNA-derived biodiversity estimates. Hill numbers are a compatible statistical framework that integrates diversity concepts by considering incidence and abundance data. They have been widely used as metrics for eDNA-based biodiversity calculation because biodiversity measurements between diversity levels or studies can be directly compared to each other ^48–51^. Based on the acquired eDNA data set, we calculated Hill numbers at each sampling site with order q = 0, 1, and 2 according to equations (1–2), which are analogue to species richness, the exponential of Shannon diversity, and the inverse of the Simpson index, respectively ^48^. For q = 1, there is a singularity problem for the equation; therefore, equation (2) was used instead.

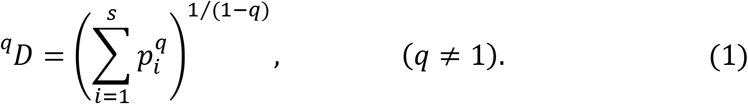

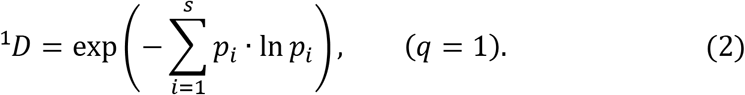

Here, *s* is the number of ZOTUs at each site, *p_i_* is the relative abundance of ZOTU *i*.

### Physiological traits in terrestrial ecosystems by Sentinel-2

We used Sentinel-2 derived measures to describe the functional diversity of the terrestrial ecosystems. We adapted a method developed by Helfenstein, 2018 ^52^, which successfully applied the terrestrial ecosystem functional diversity mapping ^22^ to Sentinel-2 MSI data, to map physiological traits at a 20-m resolution and then calculate terrestrial ecosystem functional diversity ^53^. Specifically, we used chlorophyll content (CHL), anthocyanin content (ANT), carotenoid content (CAR), and water content (WAT) to construct a four-dimensional functional space. Chlorophyll (green pigment) helps plants capture energy from light in the photosynthesis reaction; anthocyanin (blue, red, and purple pigment) replaces chlorophyll during leaf senescence process; carotenoid (orange and yellow pigment) prevents possible damage in stress conditions; water content reflects dry weight and drought stress among the plants ^25^. Hence, these traits can integrally capture the presence and conditions of vegetation ^52^.

All physiological traits were computed on Google Earth Engine (GEE), a cloud-based platform for spatial analysis ^69^. We selected Sentinel-2 MSI Level-2A calibrated surface reflectance (SR) image collections between June and August in 2017, as no SR images were produced at the time of eDNA sampling. Based on a cloud-free image acquired by employing a median filter to the selected image collections, we calculated ten indices of CHL, ANT, CAR, and WAT.

#### 1) **CHL**

the red-edge chlorophyll index (*CI_re_*, equation 3), the green chlorophyll index (*CI_g_*, equation 4), the Medium Resolution Imaging Spectrometer (MERIS) terrestrial chlorophyll index (*MTCI,* equation 5), and the normalized difference red-edge 1 and 2 (*NDRE*1 and *NDRE*2, equations 6–7).

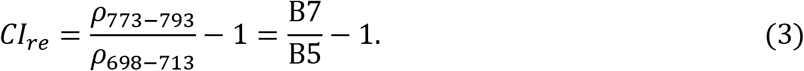

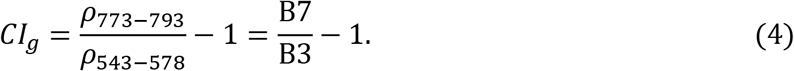

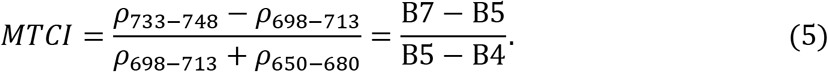

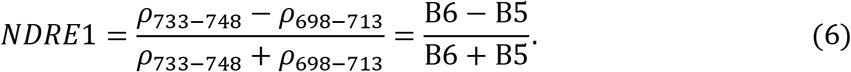

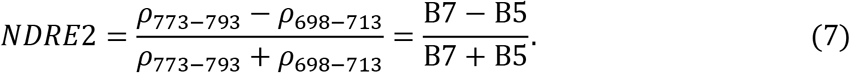

#### 2) ANT

the anthocyanin reflectance index 1 and 2 (ARI*1* and *ARI*2, equations 8–9), and the red-green ratio (*RGR,* equation 10).

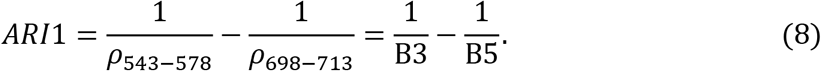

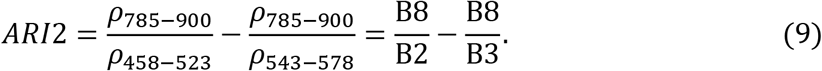

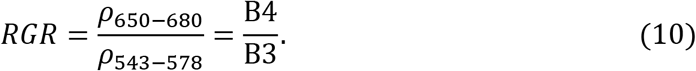

#### 3) CAR

the carotenoid reflectance index 1 (*CRI*1, 11), and the plant senescence reflectance index (*PSRI,* equation 12).

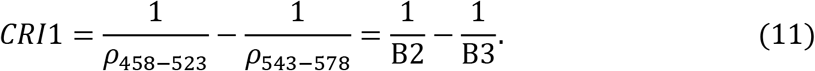

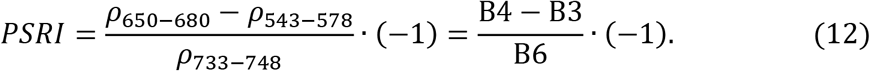

#### 4) WAT

the normalized difference infrared index (*NDII,* equation 13).

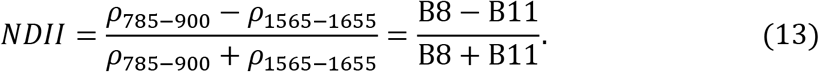

*ρ_xxx-xxx_* and B*X* represent the band of Sentinel-2 MSI.

To remove urban and water areas, we calculated the normalized difference vegetation index (*NDVI,* equation 14), and then masked out the non-vegetated pixels by setting a criterion of NDVI < 0.4.

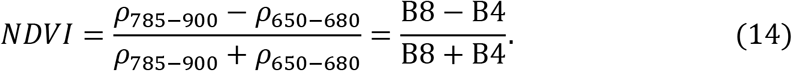

All the calculated indices were re-projected to the CH1903 projection.

### Selection of physiological traits

To reduce collinearity, we chose one trait in each physiological trait dimension. We computed a correlation matrix of all the normalized physiological traits (Fig. S8a) and enumerated all possible four-trait subsets. For each subset, we calculated the Frobenius norm (||***A***||_*F*_) of the correlation matrix (***A***), according to equation (15).

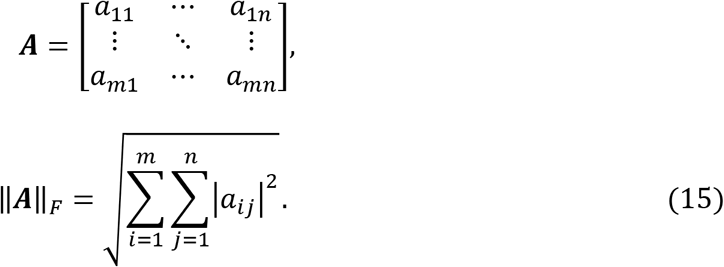

Next, we found the optimal subset with the least Frobenius norm (Fig. S8b). The selected traits were *CI_re_* (CHL), *ARI*1 (ANT), *PSRI* (CAR), and *NDII* (WAT). We observed less collinearity among the selected traits except for CHL against WAT, where positive correlations are unavoidable because the process of photosynthesis is tightly linked to chlorophyll and water availability (Fig S9).

### Catchment data and distance buffers

We used the digital elevation model (DEM) of the study area provided by the Swiss Federal Institute of Topography (Swisstopo) to extract the catchment of each eDNA sampling site. ArcGIS software (version 10.3) was used to generate a flow direction map based on the DEM. We produced a catchment map with flow distance for each site by tracing the water flow direction of each pixel and recording its flow distance to the site. Distance buffers of each sampling site were created by setting the spatial interval to 0.05 km for 0–10 km and 0.1 km for 10–20 km.

### Terrestrial ecosystem functional diversity across distance

We chose functional divergence (FDiv) among three types of functional diversity (functional richness, functional divergence, and functional evenness) because FDiv best captured the variation of terrestrial ecosystem functions and was the most robust to noises and outliers ^22^. For each sampling site with a distance buffer, based on the normalized selected traits, we extracted four-dimensional trait value vectors (*V_i_*) from vegetated pixels (i = 1,2, …, s) that covered by the distance buffer and calculated FDiv by following equations (16–19).

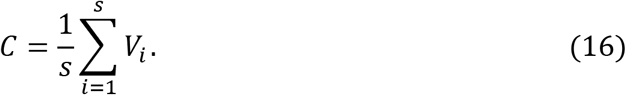

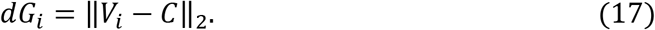

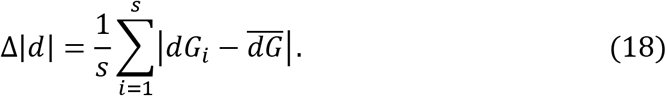

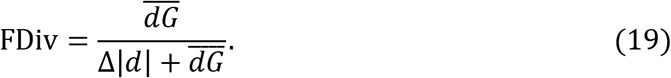

*s* is the number of vegetated pixels in the distance buffer; *C* is the center of gravity of all vectors; *dG_i_* is the Euclidean distance between the vector of *i^th^* pixel (*V_i_*) and the center of gravity (*C*). 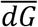 is the mean Euclidean distance of all vectors to the center of gravity (*C*).

### Linear regression model and uncertainty estimation

Due to uncertainties in both eDNA and RS measurements, we used a model II simple linear regression model to evaluate the correlation between Hill numbers and FDiv of surrounding terrestrial ecosystems across distance, using R^2^ as a metric. As distance increased, sampling sites were removed from the regression model if their catchments were already entirely covered by distance buffer (Fig. S10). To estimate uncertainties, we adopted a bootstrap framework by subsampling 70% of the available sampling sites 1,000 times, and then calculated the standard deviation of the bootstrapped R^2^ results.

### Null models for comparison

We developed null models to ensure that the spatial association between aquatic and terrestrial ecosystems was not a measurement artifact. Specifically, the spatial location of pixels (with their respective functional diversity measurement) within the river catchment were randomly shuffled in space 1,000 times, followed by calculating FDiv for each sampling site according to the same distance buffers generated before. Then, model II simple linear regression was performed to evaluate the correlation between the eDNA data and the shuffled RS data. We observed gradually increasing curves across Hill number orders without peaking signals (Fig. 5 and S11). These evidenced that the spatial fingerprint of biodiversity was a true signal from the spatial layout of the terrestrial ecosystem functional diversity, and was not an artifact.

### Evaluation of contributions of vegetation productivity and terrestrial ecosystem functional diversity

We calculated the enhanced vegetation index (EVI, equation 20), which can be used to estimate vegetation productivity ^58,70^. The EVI values were averaged across the distance buffers after excluding non-vegetated pixels.

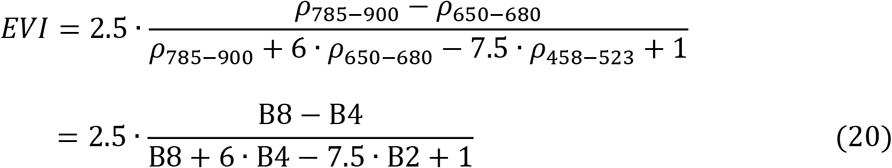

Then, we used linear models summarized in ANOVA tables with sequential (type I) tests to evaluate the relative contributions of EVI and FDiv to the Hill numbers (Hill) across distance, by equations (21–22).

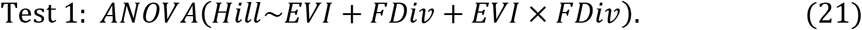

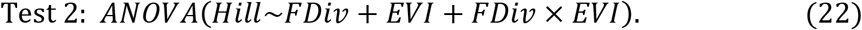

*EVI* × *FDiv* and *FDiv* × *EVI* were interaction terms. The relative contributions of EVI and FDiv are shown in Fig. S5.

## Acknowledgements

We thank Chelsea Little for support during fieldwork, Luca Carraro for help extracting catchment information, and Isabelle Helfenstein and Enrico Bertuzzo for their help with functional divergence computation. F.A. is funded by the Swiss National Science Foundation Grants No 31003A_173074 and PP00P3_179089, and F.A, F.M., and M.S. by the University of Zurich Research Priority Programme on Global Change and Biodiversity (URPP GCB).

