## Supplementary Material for "A spatial fingerprint of land-water linkage of biodiversity uncovered by remote sensing and environmental DNA"

#### Supplementary Materials

**Figures S1 – S11**

**Table S1 – S4**

### Figures

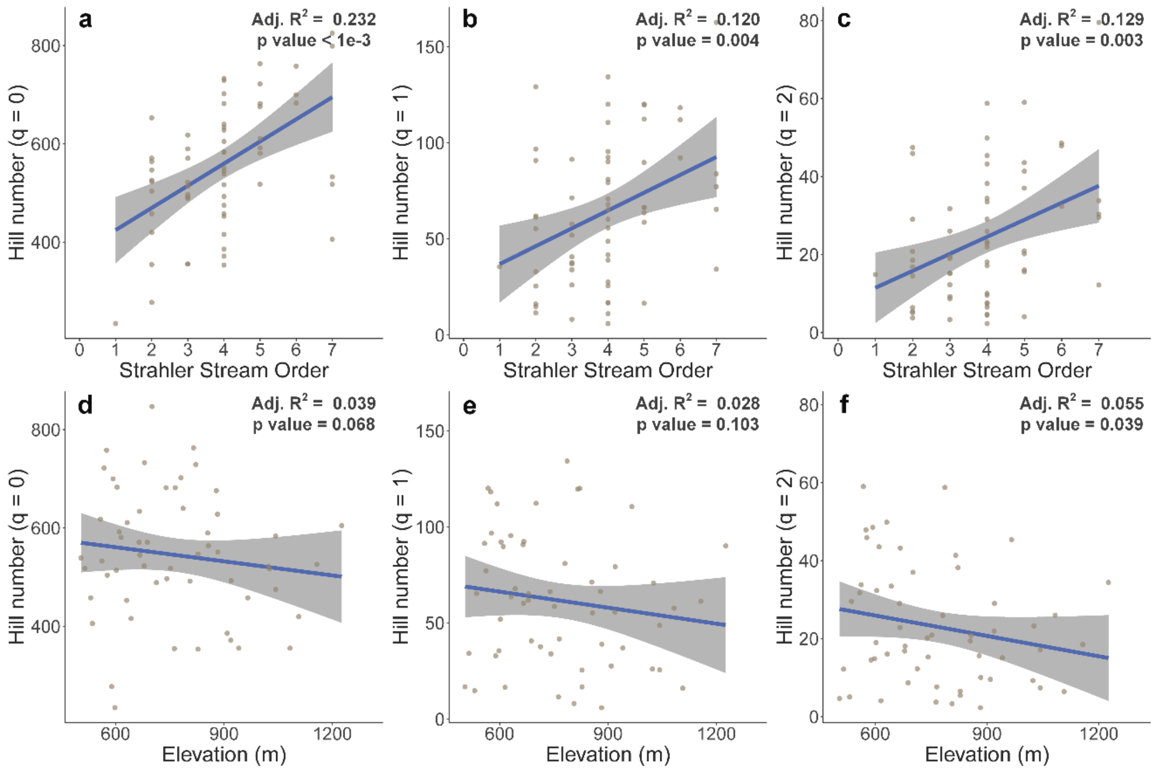


**Fig. S1** The correlations between eDNA-derived biodiversity and Strahler stream order and elevation, respectively, at sampling sites: **a** Hill number with order q = 0 against Strahler stream order; **b** Hill number with order q = 1 against Strahler stream order; **c** Hill number with order q = 2 against Strahler stream order; **d** Hill number with order q = 0 against elevation; **e** Hill number with order q = 1 against elevation; **f** Hill number with order q = 2 against elevation. The Adj.$R^2$ and p-value were estimated using an ordinary linear regression model.

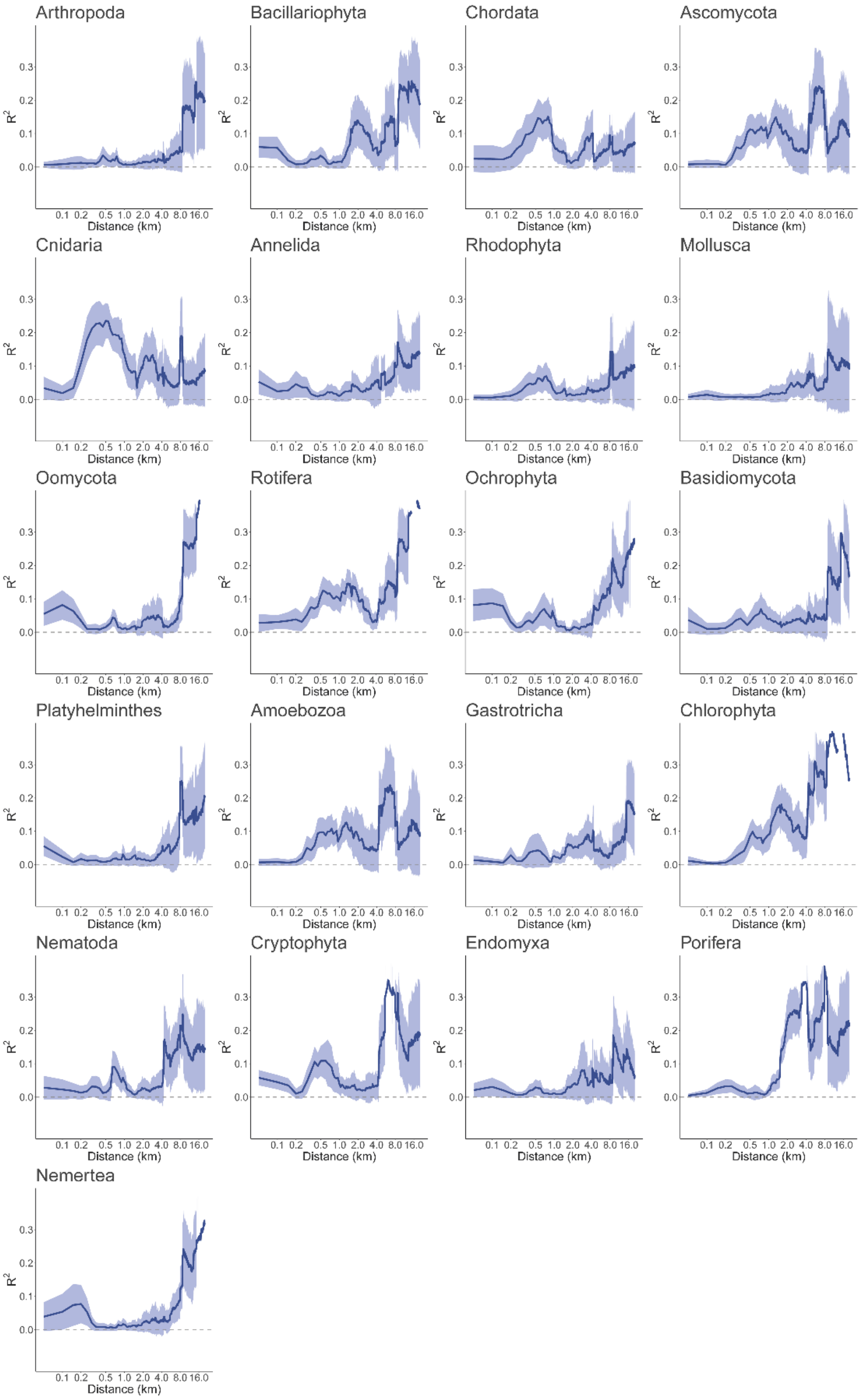


**Fig. S2** The $R^2$ of linear regressions (± standard deviation) between the relative abundance of major taxonomic groups at the phylum level and remote sensing (RS)-based terrestrial ecosystem functional divergence (FDiv) across distance.

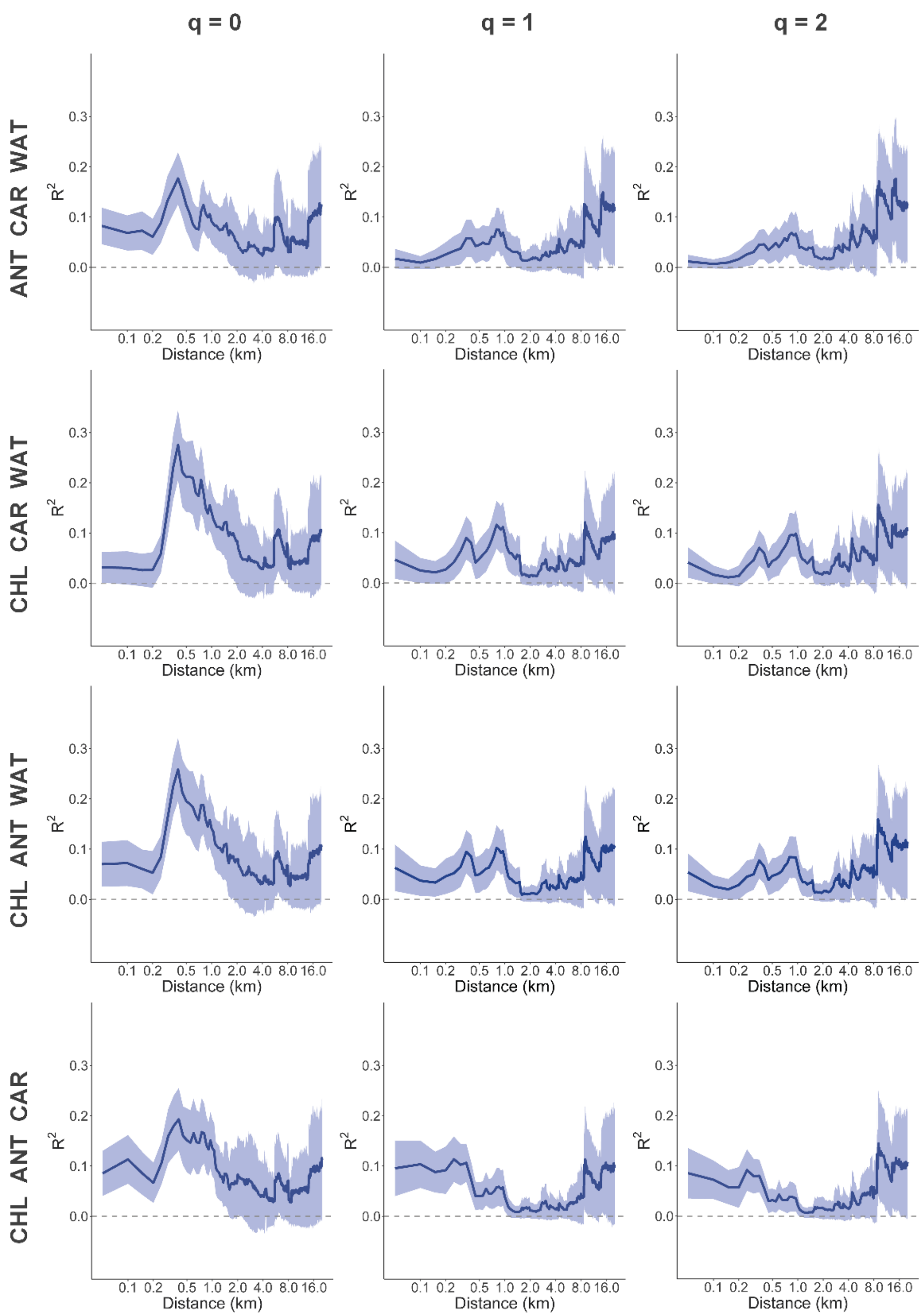


**Fig. S3** The $R^2$ of linear regression (± standard deviation) between eDNA-derived biodiversity (Hill numbers) with Hill number orders q = 0, 1, 2, and RS-based FDiv on three out of the four physiological trait dimensions (CHL: chlorophyll content, ANT: anthocyanin content, CAR: carotenoid content, WAT: water content) across distance. The calculations were done in a leave-one-out approach (compared to the main analysis, in which all four were included).

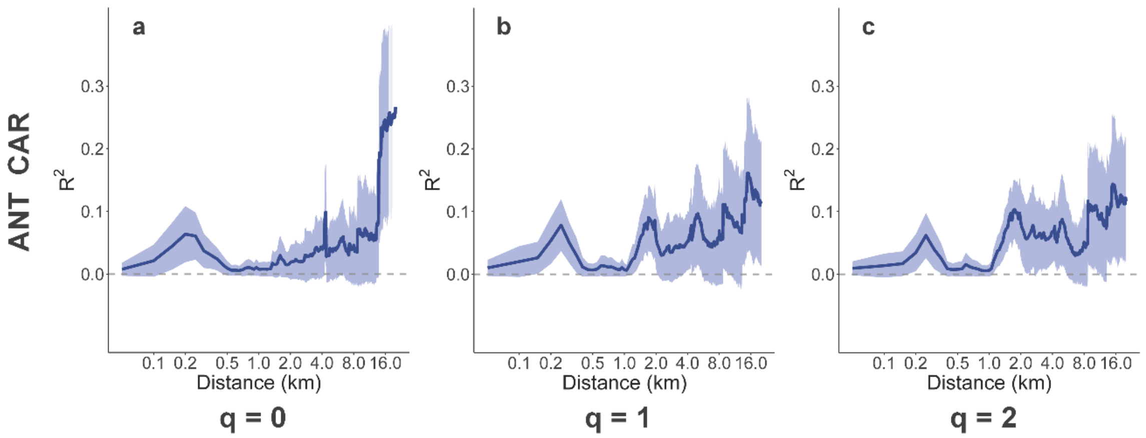


**Fig. S4** The $R^2$ of linear regression (± standard deviation) between eDNA-derived biodiversity (Hill numbers) with Hill number orders **a** q = 0, **b** q = 1, **c** q = 2, and RS-based FDiv on two physiological trait dimensions (ANT: anthocyanin content, CAR: carotenoid content) across distance.

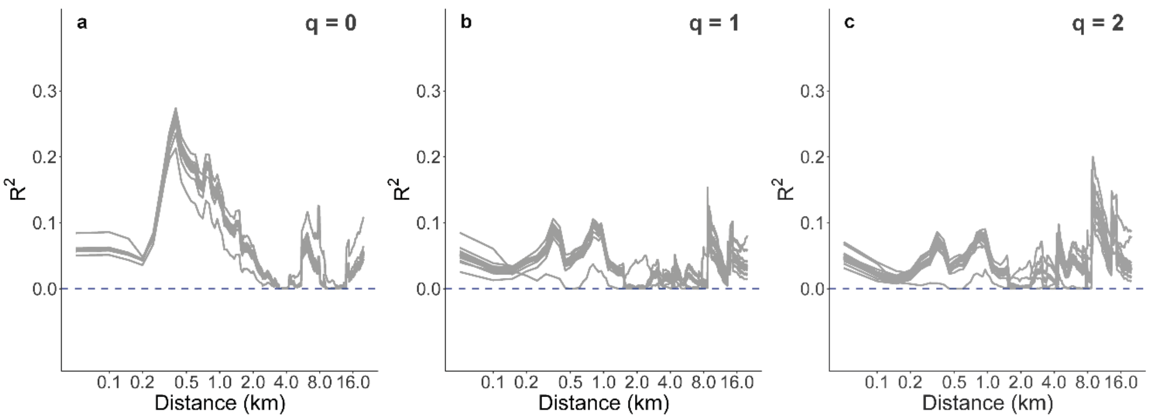


**Fig. S5** The $R^2$ of linear regressions between eDNA-derived biodiversity (Hill numbers) with Hill number orders **a** q = 0, **b** q = 1, **c** q = 2 by removing each major taxonomic group and RS-based FDiv across distance. The calculations were done in a leave-one-out approach. Each grey line represents the $R^2$ across distance after removing one taxonomic group at the phylum level from the eDNA data.

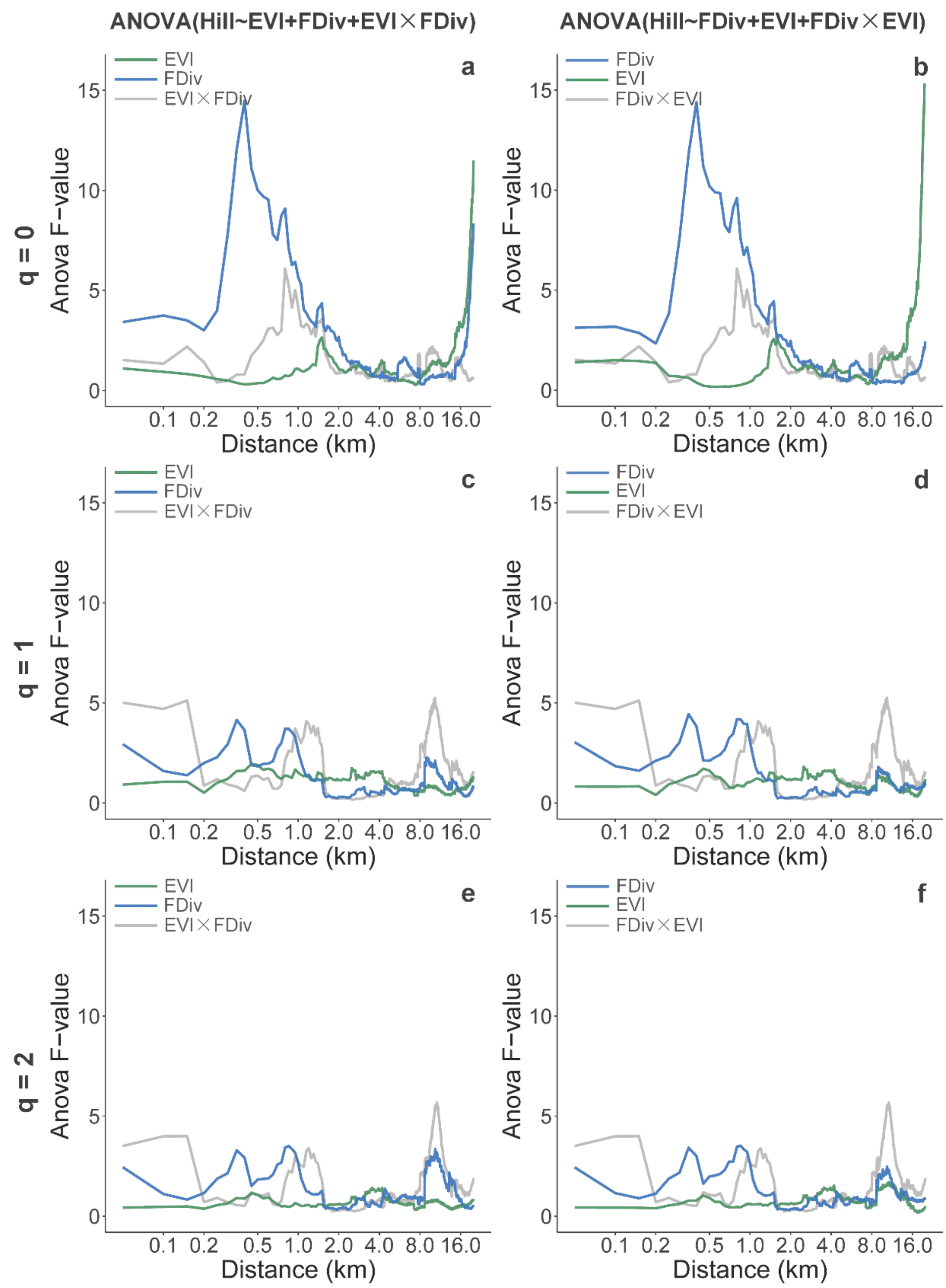


**Fig. S6** The F-values of the enhanced vegetation index (EVI), FDiv, and their interaction term in type I ANOVA tests (test 1 and test 2) with Hill number in orders q = 0, 1, 2 across distance.

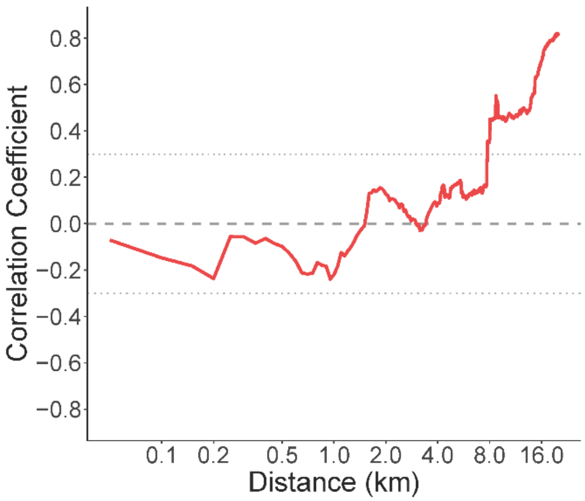


**Fig. S7** Correlation coefficients between EVI and FDiv across distance.

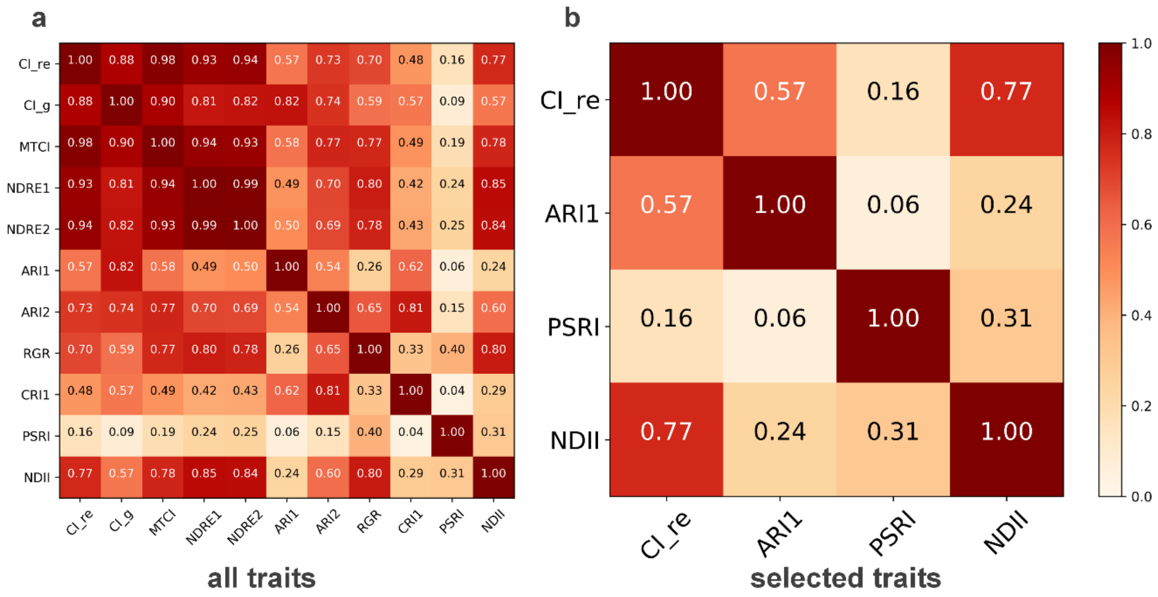


**Fig. S8** Correlation matrices of **a** all physiological traits available and **b** the selected traits used (CHL: the red-edge chlorophyll index ($CI_{re}$), ANT: the anthocyanin reflectance index 1 (*ARI1*), CAR: the plant senescence reflectance index (*PSRI*), WAT: the normalized difference infrared index (*NDII*)).

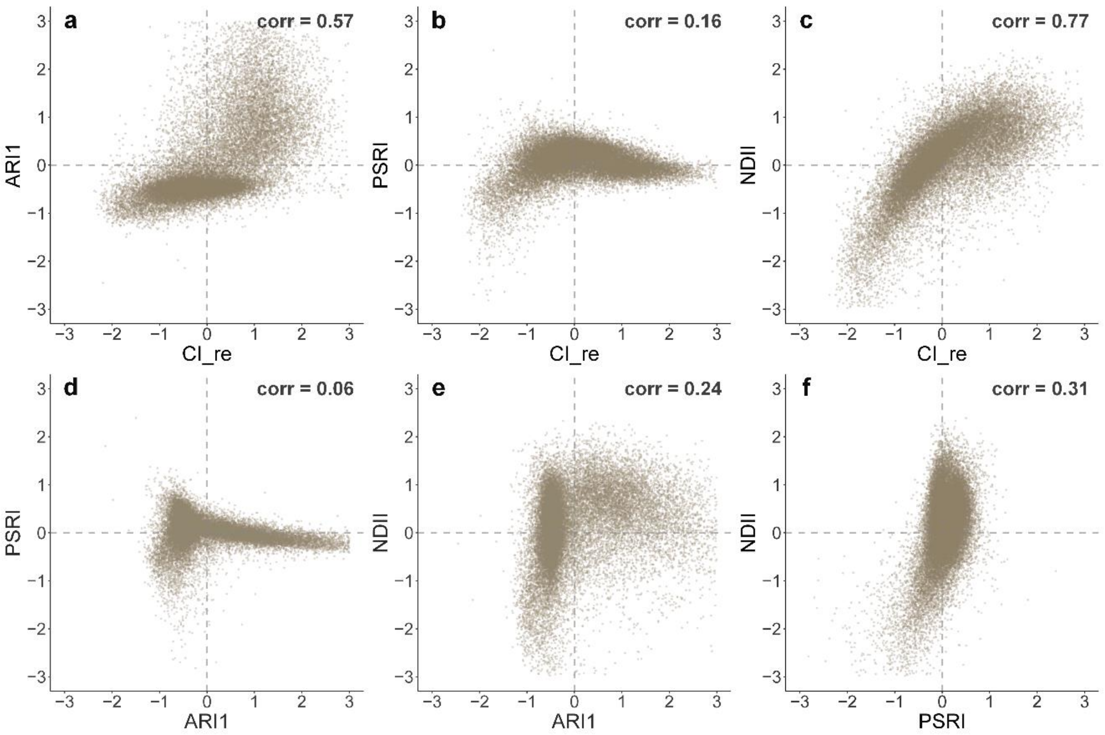


**Fig. S9** Scatter plots of the selected traits used (CHL: the red-edge chlorophyll index ($CI_{re}$), ANT: the anthocyanin reflectance index 1 (*ARI1*), CAR: the plant senescence reflectance index (*PSRI*), WAT: the normalized difference infrared index (*NDII*)): **a** *ARI1* against $CI_{re}$, **b** *PSRI* against $CI_{re}$, **c** *NDII* against $CI_{re}$, **d** *PSRI* against *ARI1*, **e** *NDII* against *ARI1*, **f** *NDII* against *PSRI*.

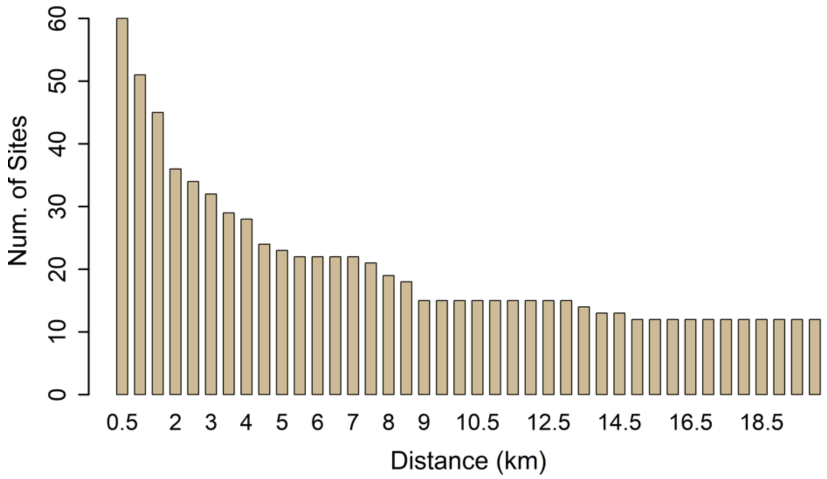


**Fig. S10** Number of sampling sites included in linear regression models across distance. Sites were removed once the distance buffer extended their entire catchment.

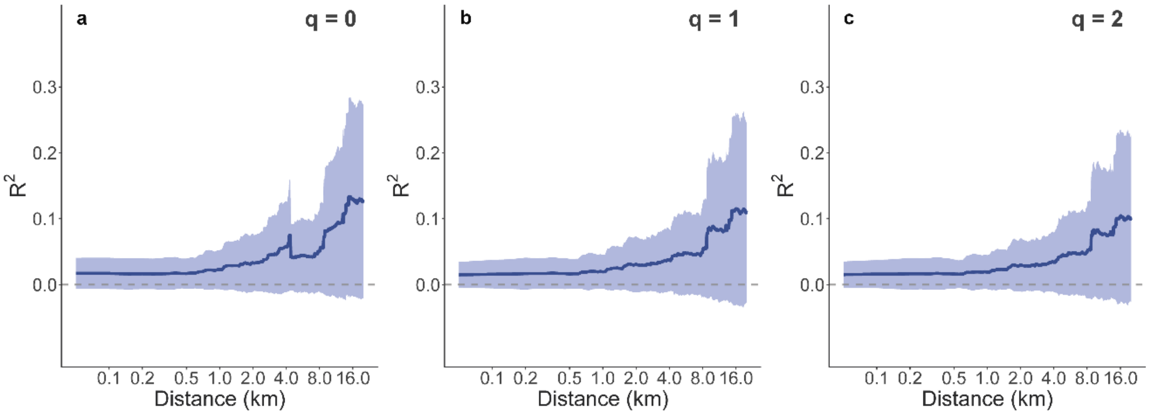


**Fig. S11** The $R^2$ of linear regression (± standard deviation) between eDNA-derived biodiversity (Hill numbers) with Hill number orders **a** q = 0, **b** q = 1, **c** q = 2, and RS-based FDiv on shuffled RS data across distance. We observed an increase of standard deviations, especially for distances > 8 km, driven mainly by the small number of sites available for calculations at these distances (see also Fig. S10). Therefore, conclusions are less robust for distances larger than about 8 km.

#### Tables

**Tab. S1** The highest $R^2$ of linear regression between eDNA-derived biodiversity (Hill numbers) with Hill number orders q = 0, 1, 2, and RS-based terrestrial ecosystem functional divergence (FDiv) on a subset of four physiological trait dimensions (CHL: chlorophyll content, ANT: anthocyanin content, CAR: carotenoid content, WAT: water content) across distance.

| Physio-traits | q = 0 | q = 1 | | q = 2 | |
|---|---|---|---|---|---|
| | 0.40 km | 0.35 km | 0.80 km | 0.35 km | 0.85 km |
| CHL ANT CAR WAT | 0.259 | 0.096 | 0.103 | 0.075 | 0.086 |
| ANT CAR WAT | 0.177 | 0.058 | 0.075 | 0.046 | 0.069 |
| CHL CAR WAT | 0.275 | 0.090 | 0.116 | 0.070 | 0.097 |
| CHL ANT WAT | 0.258 | 0.095 | 0.102 | 0.077 | 0.084 |
| CHL ANT CAR | 0.193 | 0.106 | 0.058 | 0.080 | 0.038 |
| ANT CAR | 0.024 | 0.030 | 0.010 | 0.025 | 0.006 |

**Tab. S2** Correlations of the bands between 2016 and 2017 in Sentinel-2 Multi-Spectral Instrument (MSI) data. All the calculations were based on Level-1C top of atmosphere reflectance data.

| | B2 | B3 | B4 | B5 | B6 | B7 | B8 | B11 |
|---|---|---|---|---|---|---|---|---|
| correlation coefficient | 0.845 | 0.900 | 0.870 | 0.917 | 0.911 | 0.890 | 0.896 | 0.942 |

**Tab. S3** The correlations of physiological traits between 2016 and 2017 based on Sentinel-2 Level-1C data.

| | CHL ($CI_{re}$) | ANT (ARI1) | CAR (PSRI) | WAT (NDII) |
|---|---|---|---|---|
| correlation coefficient | 0.741 | 0.795 | 0.773 | 0.827 |

**Tab. S4** The mean temperature (T) and precipitation (P) in the summer (Jun–Aug) of 2016 and 2017 at three measuring stations near the river catchment.
(Data source: https://www.meteoswiss.admin.ch/home/climate/swiss-climate-in-detail/homogeneous-data-series-since-1864.html)

| | T in 2016 (°C) | T in 2017 (°C) | T normal (°C) | P in 2016 (mm) | P in 2017 (mm) | P normal (mm) |
|---|---|---|---|---|---|---|
| Zürich/Fluntern | 18.2 | 19.5 | 17.7 | 116.3 | 119.9 | 125.3 |
| St. Gallen | 16.9 | 18.1 | 16.3 | 164.8 | 161.4 | 177.3 |
| Säntis | 6.0 | 7.4 | 5.3 | 285.3 | 286.6 | 264.7 |
